# Morphomolecular Longitudinal Endoscopy of Carcinogenesis

**DOI:** 10.1101/2025.08.15.665660

**Authors:** Jianrong Qiu, Priyanka G. Bhosale, Vishal Kumar, Jeyrroy Gabriel, Shiyue Liu, Ciro Chiappini, Richard J. Cook, Mads S. Bergholt

## Abstract

Current diagnostic approaches for head and neck cancer rely primarily on white-light endoscopy and tissue biopsy, providing only static, episodic assessments that fail to capture dynamic changes in tumour biology over time. This absence of continuous, detailed molecular and microscopic structural information makes early identification of dysplasia or cancer difficult and can delay critical treatment. Here, we introduce a unified, label-free endoscopic platform that combines Raman spectroscopy (RS) and optical coherence tomography (OCT) within a miniaturised, forward-viewing fibre-optic probe. This integration enables simultaneous video-rate OCT imaging and subsecond RS acquisition linking morphological and molecular tissue trajectories *in vivo*. We developed an explainable AI-based fusion framework that learns joint temporal representations of biochemical and structural features to classify tissue states while highlighting the most influential predictors. In a longitudinal 4NQO-induced murine model of carcinogenesis, the system tracked epithelial transformation in real-time from hyperplasia through dysplasia to neoplasia. By capturing temporally resolved morphomolecular trajectories *in vivo*, this platform establishes a new paradigm for early cancer detection and real-time surveillance, supporting timely and individualised clinical decision-making.

## Introduction

Head and neck squamous cell carcinoma (HNSCC) remains a global health challenge, imposing substantial health, social and economic burdens^1^. Despite advances in treatment, the prognosis for HNSCC remains poor, as majority of cases are diagnosed at advanced stages. This makes early detection crucial, as it creates a window of opportunity for timely disease management, better clinical outcomes, and the preservation of vital functions such as speech and swallowing, ultimately enhancing patients’ quality of life. Some precancerous lesions remain latent without ever progressing, while others (10-25%) are more likely to advance to malignancy or develop resistance to treatment^2-4^. Leaving high-risk lesions *in situ* leads to evolution of the tumour microenvironment and exacerbation of disease, with devastating consequences for the patient. Therefore, monitoring the disease trajectory of lesions is of significant clinical importance to initiate targeted and aggressive treatment (e.g., resection, ablation or neoadjuvant chemotherapy etc.). Conventional diagnostic techniques are based on visual examination, white light or narrowband endoscopy. When combined with biopsy and histopathology this is inherently invasive and hinder continuous longitudinal monitoring due to the need for repeated tissue sampling.

HNSCC is a multistep disease driven by alterations in genomic, cellular, molecular, and tissue microarchitecture^5^. The diverse oncogenomic landscape largely stems from the selection and clonal expansion of tumour-initiating cells that acquire various molecular and genetic alterations^6^. This includes frequent mutations of tumour suppressor genes such as TP53, FAT1, NOTCH1, PIK3CA and CASP8. Many of these mutations disrupt cellular processes that regulate proliferation and differentiation, resulting in a progressively altered molecular profile within the tissue^7^. Concurrently, these cellular and molecular changes are accompanied by microscopic morphological transformations. Dysplastic tissues often exhibit loss of epithelial cohesion, disrupted stratification, basal cell hyperplasia, epithelial thickening, and irregularly shaped rete pegs. These structural abnormalities are strongly associated with malignant transformation and increased risk of lesion recurrence^8^. As dysplasia advances to carcinoma, tumour-initiating cells invade the surrounding stroma, breaking down normal structural boundaries and leading to highly disorganised tissue architecture.

Fibre-based optical imaging techniques have emerged as promising tools, offering a unique combination of label-free, non-invasive modalities that deliver high-resolution morphological imaging alongside molecular analysis, paving the way for more dynamic and informative tissue assessment. Techniques such as Raman spectroscopy (RS)^9-11^, optical coherence tomography (OCT), confocal laser endomicroscopy (CLE)^12^, and multiphoton microscopy^13-15^ including second harmonic generation (SHG), two-photon excited fluorescence (TPEF), and coherent anti-Stokes Raman spectroscopy (CARS), have demonstrated considerable potential for cancer diagnostics. RS exploits the inelastic scattering of light to provide detailed vibrational signatures of molecular bonds, thereby enabling the identification of biomolecular fingerprints and disease markers. RS provides rich molecular fingerprints of tissue pathology, yet is blind to the morphological context in which these changes occur. OCT complements this gap by offering high-resolution, cross-sectional reconstructions of tissue microarchitecture, enabling the visualisation of subtle structural perturbations of the disease onset and progression^16^.

Integrating RS and OCT into a single, miniaturised endoscopic probe presents major engineering challenges. RS depends on efficient photon collection and distal optical filtering. Meanwhile, OCT relies on beam scanning and integrated focusing optics to generate high-resolution images. These fundamentally different optical demands create a complex set of design trade-offs, making it difficult to merge both modalities into one compact fibre-optic probe compatible with endoscope instrument channels. Another critical challenge is balancing a sufficient optical field of view (FOV) with the strict mechanical constraints required for miniaturisation within standard endoscopic channels. Historically, pioneering efforts to combine RS and OCT in a fibre-optic format have therefore been limited to large or side-viewing designs with dimensions that exceed requirements needed in clinical endoscopy^17-19^. Temporal co-registration remains another challenge: RS acquires spectra on subsecond timescales, whereas OCT provides video-rate images. This temporal discrepancy demands novel fusion strategies that can integrate the disparate data into a joined, coherent representation.

Here, we introduce a unified, forward-viewing endoscopic platform that integrates Raman spectroscopy and optical coherence tomography (RS-OCT). The system incorporates a miniaturized, high throughput fibre-optic probe with a spliced-fibre architecture and distal beam scanning. Designed to be compatible with standard clinical endoscopes, we applied this probe in a longitudinal *in vivo* mouse model of H&N carcinogenesis. Our approach enables high-resolution, video-rate longitudinal OCT imaging of tissue microarchitecture alongside subsecond RS-based molecular profiling. To interpret the multimodal data, we developed a quantitative analysis pipeline and an explainable deep learning-based fusion framework that classifies tissue by integrating biomolecular and morphological signatures. Collectively, this technology provides a powerful, non-invasive strategy for comprehensive temporal tissue characterisation in internal organs, with strong translational potential to inform precision diagnostics and guide personalised therapeutic decision-making in oncology.

## Results

### Unified RS-OCT fibre-optic endoscopy

We developed a compact, integrated endoscopic platform that unites RS and OCT within single, miniaturised fibre-optic probes (2.2 mm and 3.4 mm diameter versions; Fig. 1a,b). RS was implemented with 785 nm laser excitation and high-throughput signal collection across the 800-1800 cm^−1^ fingerprint window, while OCT imaging employed a custom-built ∼1300 nm spectral-domain system. The developed miniaturised probe utilised a sandwich-like fibre configuration: two arrays of RS excitation/collection fibres were angled toward the optical axis (Supplementary Fig. S1), flanking a centrally positioned single-mode fibre for OCT (Fig. 1d). Since RS is not compatible with distal micro lenses, the OCT light focusing was achieved by splicing the single mode fibre with a no core fibres (NCFs) and a gradient index fibre (GIF) maintaining focusing of ∼0.5 mm in tissue (Fig. 1e). This ensures that no parasitic Raman signals were generated from any distal optics. This design enabled OCT B-scanning across a central opening, providing a 1 mm field of view with axial and transverse resolutions of 10 μm and ∼25 μm, respectively. By maintaining a consistent spatial relationship between the structural and molecular sensing components, this symmetric design allows for precise co-localisation of morphological features with their corresponding biomolecular signatures. The internal structure of the fibre probe was 3D printed to house a piezo tube enclosed by biocompatible stainless steel and sealed with a 10-degree tilted CaF_2_ window to reduce OCT back reflections (Fig. 1f).

**Fig. 1:**
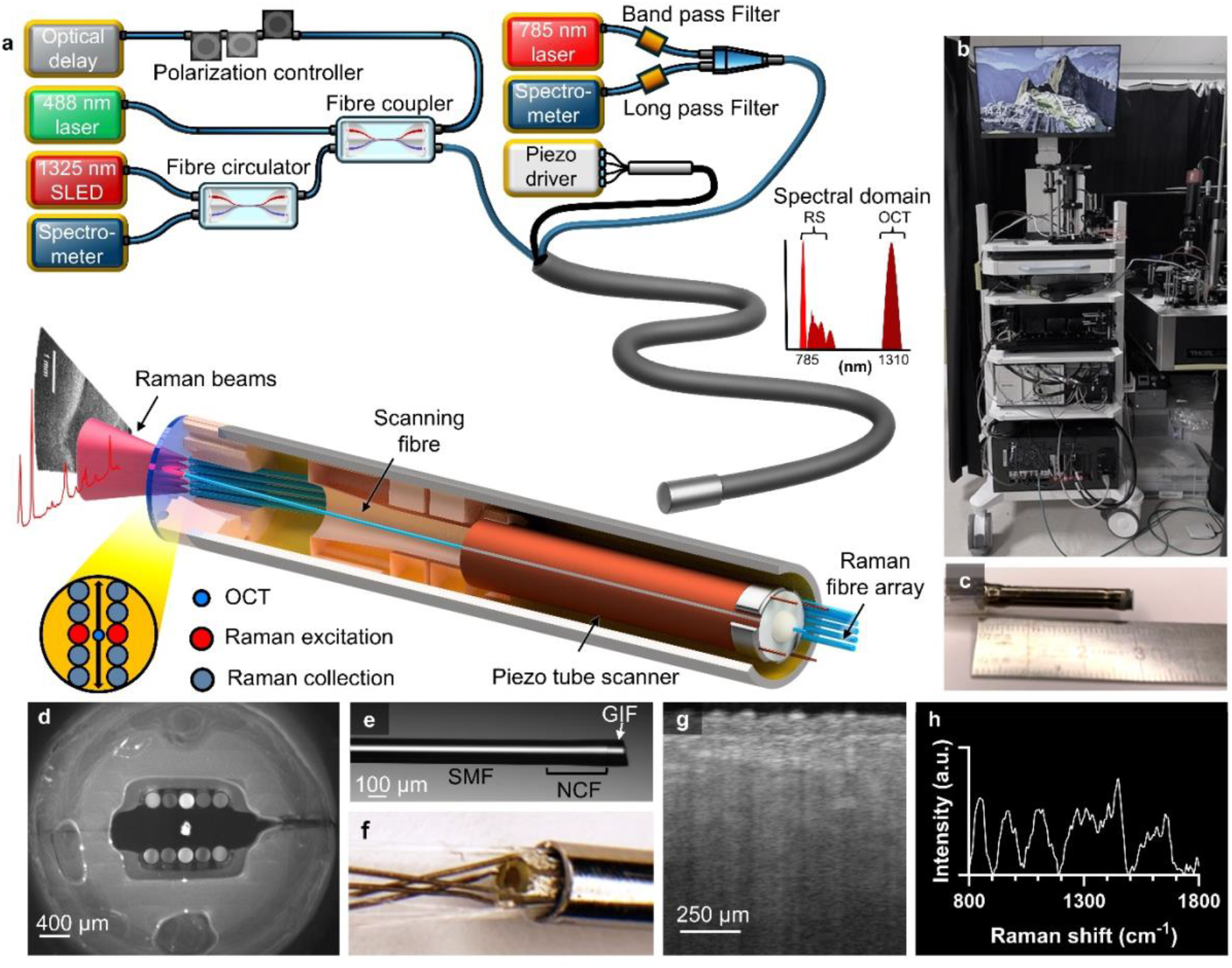
Unified multimodal fibre-optic probe for Raman spectroscopy and OCT. **a**, The multimodal Raman-optical coherence tomography (RS-OCT) fibre-optic probe, designed based on a spectral-domain OCT scanning platform, utilises a compact 2.2 or 3.4 mm piezoelectric (PZT) tube actuator for beam scanning. The Raman spectroscopy (RS) component operates with 785 nm laser excitation and incorporates a high-throughput spectrograph for efficient signal detection. **b**, Complete assembly of the RS-OCT system mounted on a medical-grade trolley. **c**, Photograph of the developed multimodal RS-OCT fibre probe. **d**, En face view of the fibre probe, showing the central OCT fibre surrounded by two linear arrays of excitation/collection fibres dedicated to Raman signal acquisition. **e**, Detailed structure of the spliced single-mode fibre (SMF) used in the OCT path, consisting of a segment of SMF for light delivery, followed by a no-core fibre (NCF) for beam expansion, a gradient-index fibre (GIF) for focusing, terminated with an angle-cleaved surface for back-reflection suppression. **f**, assembled probe showing the fibre bundles and electrical wiring. **g**, Representative *in vivo* OCT imaging demonstrating high-resolution structural visualisation. **h**, Corresponding *in vivo* Raman spectra acquired from the same region, providing complementary molecular information.

Processing real-time multimodal datasets acquired at different timescales remains a major challenge. We constructed a parallel data acquisition and processing framework (Supplementary Fig. S2) implemented with CPU based RS analysis and GPU based OCT image processing. The OCT images were reconstructed using background removal, k-domain linearisation, Fast Fourier Transform (FFT) and offset/scaling. Raman spectra were processed using background subtraction, autofluorescence removal using a polynomial background subtraction and vector normalisation. This integrated framework allows us to measure real-time colocalised high quality OCT images (Fig. 1g) and tissue Raman spectra (Fig. 1h). Temporal coregistration of RS and OCT data was achieved by synchronising each one-second RS acquisition with a randomly selected frame from the corresponding OCT sequence, acquired at 46 frames per second. An OCT frame within the RS acquisition window was randomly selected to avoid consistently sampling the same phase of endoscope-induced motion, ensuring that the co-registered morphological and molecular data represented the same tissue state in time.

### *In vivo* imaging in a murine model of head and neck carcinogenesis

In a murine model of head and neck carcinogenesis, we demonstrated real-time, *in vivo* imaging, providing a powerful view of the disease process at both molecular and microstructural levels. Oral tissue was used as test bed to represent an inherently challenging multiphenotypic tissue environment. We employed a 4-Nitroquinoline N-oxide (4NQO) oral carcinogenesis mouse model, which is well established for studying disease progression and exhibits molecular and histological changes similar to those seen in human HNSCC^7^. RS-OCT data were collected longitudinally at predefined anatomical locations in both the 4NQO-treated group (n=10) and control group (n=10) at multiple time points (weeks 10, 12, 14, 16, 18, 20 and 22) (Fig. 2a & Supplementary video). At the study endpoint (week 22), visual inspection revealed lesions at multiple sites on the tongues of treated mice (Supplementary Fig. S3-S4). Histological evaluation of these tissues was then performed as the gold standard to determine the ground truth disease outcome (Fig. 2b, Supplementary Fig. S5). The lesions were histologically categorised as hyperplasia, various grade of dysplasia, squamoproliferative lesions (SPL) resembling carcinoma in situ, papilloma and invasive OSCC (Fig. 2b). Exposure to 4NQO resulted in the development of multifocal hyperplastic and/or mild dysplastic lesions in 100% of treated mice, with each mouse presenting with at least four lesions (Supplementary Fig S3). Of these, 70–80% progressed to moderate-to-severe dysplasia or squamous papillary lesions (SPL), while 40–60% developed microinvasive or invasive OSCC within 22 weeks. OCT imaging enabled clear visualisation of epithelial architecture, with control tissues displaying well-defined, stratified epithelium (Fig. 2c). We observed progressive epithelial thickening in carcinogen-treated tissues, along with increasing architectural disarray, a hallmark of dysplastic and malignant progression as confirmed by endpoint histopathology (Fig. 2b). Co-registered high-fidelity Raman spectra exhibited distinct biochemical signatures, including prominent peaks at 1301 cm^−1^ (CH_2_ vibrations from lipids/proteins), 1450 cm^−1^ (CH_2_/CH_3_ bending), and 1650 cm^−1^ (Amide I, C=O stretching), aligning with recognised spectral markers of biological tissue (Fig. 2d)^9,20^. RS revealed spectral changes between pathologies, as visualised in average difference spectra ±1 standard deviation (SD) (Supplementary Fig. S6), reflecting underlying biochemical remodeling across disease stages. These RS-OCT results establish the integrated system as a label-free, multimodal endoscopic platform for tracking carcinogenesis with both spatial and molecular fidelity

**Fig. 2:**
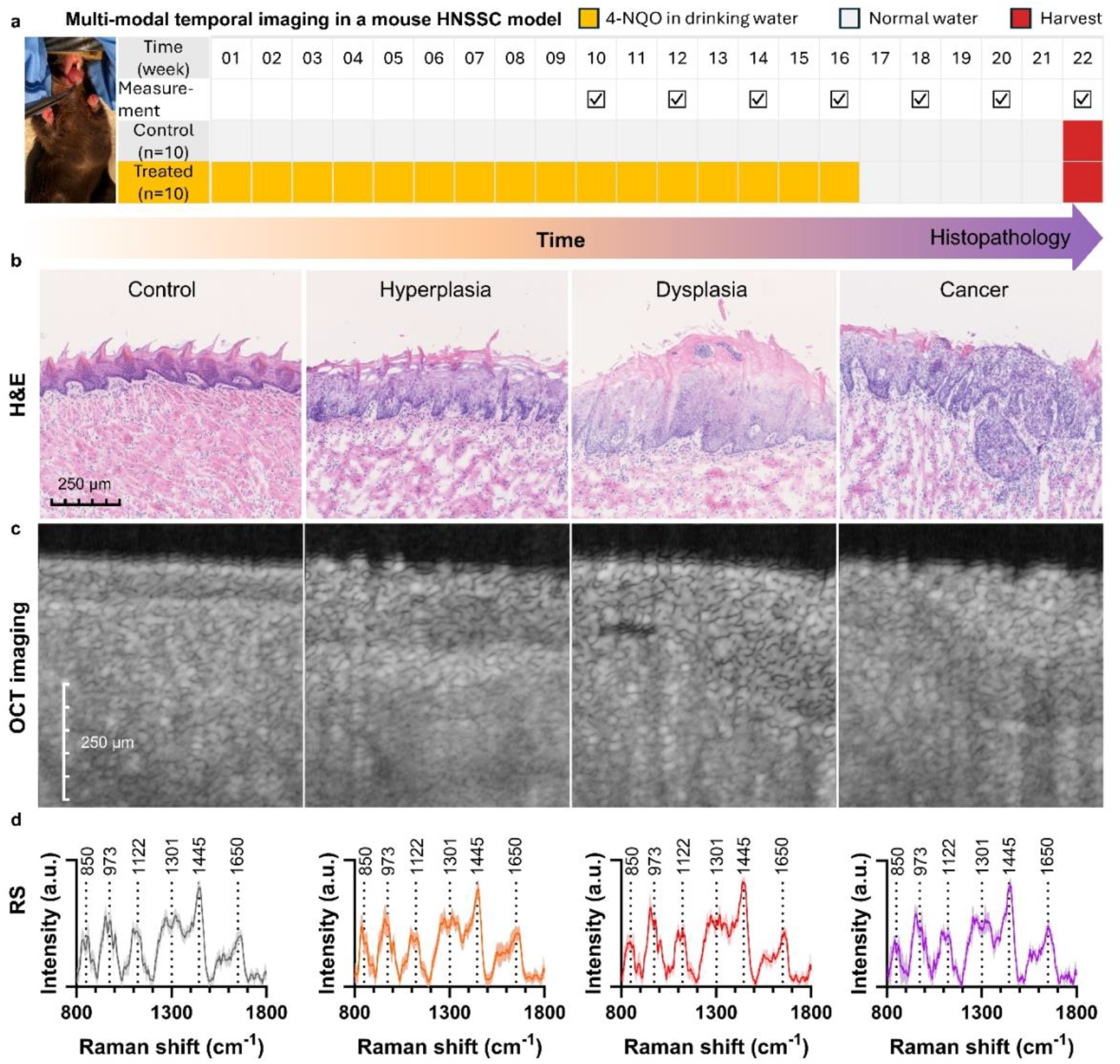
Morphomolecular *in vivo* monitoring mouse model of H&N carcinogenesis. **a**, Study design: Head and neck carcinogenesis was induced using the chemical carcinogen 4-Nitroquinoline N-oxide (4NQO). A cohort of mice (n = 10) received 4NQO diluted in drinking water at a concentration of 100 μg/mL for a 16-week treatment period. Control mice (n = 10) received regular water. After the treatment phase, all animals were returned to normal drinking water and monitored until the experimental endpoint at week 22 or until a weight loss exceeding 15% of their peak body weight was observed. RS-OCT measurements were performed longitudinally at weeks 10, 12, 14, 16, 18, 20, and 22. **b**, Representative haematoxylin and eosin (H&E) stained sections from normal, treated, dysplastic, and cancerous tissues, validating structural changes observed with OCT. **c**, *In vivo* OCT imaging at multiple time points, illustrating the progression from normal epithelium, hyperplasia to dysplasia and cancer. Imaging captures progressive epithelial thickening, heightened scattering, and increasing tissue disorganisation during carcinogenesis. **d**, Mean *in vivo* Raman spectra ±1 standard deviation (SD) corresponding to normal, hyperplasia, dysplastic, and cancerous tissue, demonstrating molecular alterations throughout disease progression.

### Longitudinal quantitative profiling of head and neck carcinogenesis

We developed an analytical framework to quantify morphological and molecular changes from multimodal data, with OCT image analysis measuring epithelial thickness and grading stratification and architectural disorganisation (Fig. 3a). Hyperplastic lesions and lesions with histologically confirmed dysplasia and to invasive OSCC exhibited progressive epithelial thickening as compared to control tissue (Fig. 3b), consistent with histopathological grading (Fig. 2b). Both dysplastic and malignant tissues showed progressive architectural disorganisation as the primary morphological indicator (Fig. 3c). This could be a result of loss of basal cell polarity together with a breakdown in the orderly maturation and differentiation of epithelial cells, a hallmark of neoplastic transformation. Crucially, our platform captured these dynamic changes noninvasively and without labels. Longitudinal analysis in a single mouse revealed continuous remodeling of epithelial architecture, with quantifiable changes in thickness and stratification over time (Fig. 3d), demonstrating the system’s capability for real-time monitoring of disease trajectory.

**Fig. 3:**
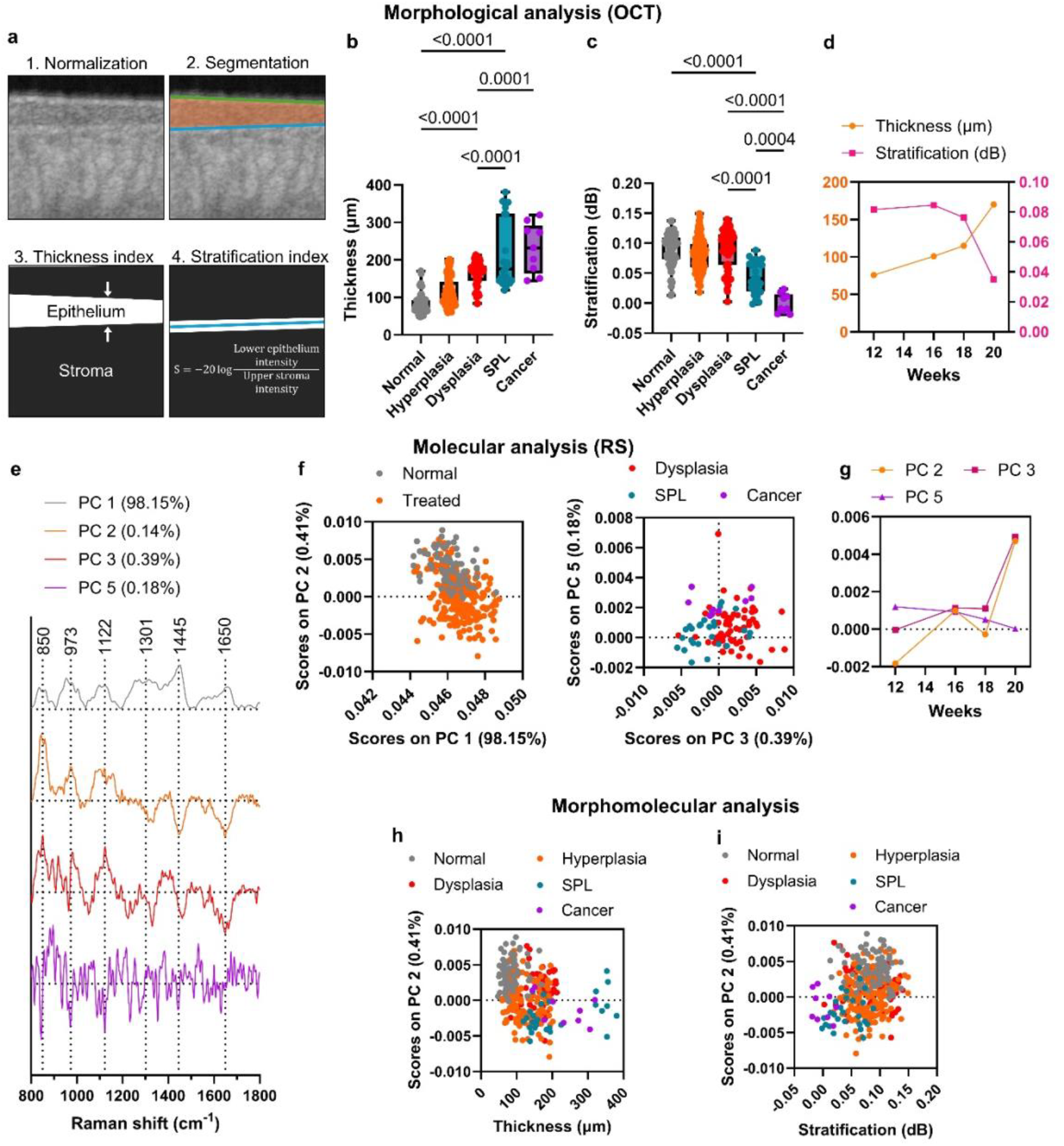
Longitudinal quantitative profiling of head and neck carcinogenesis. **a**, Quantitative workflow for optical coherence tomography (OCT) analysis, including epithelial thickness measurement and tissue stratification index calculation. **b**, OCT-derived epithelial thickness measurements for control, hyperplasia, and histologically confirmed squamous papillary lesions (SPL), dysplasia and cancer groups. **c**. OCT-based tissue stratification index for control, hyperplasia, SPL, dysplasia, and cancer cohorts. **d**, Longitudinal monitoring of epithelial thickness and stratification in a single mouse. **e**, Principal component analysis (PCA) loadings (PC1, PC2, PC3 and PC5). PCA scores (PC1 vs PC2) and (PC3 vs PC5) from histopathological confirmed control, hyperplasia, dysplasia, SPL and cancerous tissues. **f**, Longitudinal monitoring of molecular changes in a single mouse. **g**, Morphomolecular analysis showing the relationship between PC2 and epithelial thickness across tissue states, highlighting how molecular signatures captured by RS correlate with structural changes detected by OCT. **h**, Morphomolecular analysis showing the relationship between PC2 and epithelial stratification, illustrating how molecular variation aligns with the degree of tissue organisation and differentiation.

For RS molecular characterisation, we applied principal component analysis (PCA) to Raman spectra from H&N tissues, capturing dominant biomolecular variance. PCA loadings highlighted vibrational signatures of such as 850, 1301 and 1650 cm^-1^ associated with collagen, lipids and proteins (Fig. 3e). PCA score plots showed separation between control and treated tissues, reflecting disease-associated molecular shifts (Fig. 3f–g). In a single mouse, longitudinal PCA mapping revealed a continuous trajectory in spectral space, in which successive timepoints traced a directional shift along principal component axes associated with specific biochemical changes. This gradual migration reflected coordinated biochemical remodeling of the tissue microenvironment, with the spectral trajectory serving as a biomolecular fingerprint of disease progression (Fig. 3h).

To investigate how molecular alterations relate to morphological remodeling, we correlated the PCA scores from Raman spectra with epithelial thickness and stratification metrics extracted from OCT. Plotting PC2 scores against epithelial thickness revealed clustering, in which progressive thickening was accompanied by coordinated shifts in molecular composition (Fig. 3h). Similarly, PC stratification plots demonstrated that increasing architectural layering and disorganisation tracked with distinct molecular signatures along specific PC axes (Fig. 3i). Additional analyses of disease-associated PCs across samples are presented in Supplementary Fig. S7. These relationships suggest that the same biochemical pathways captured in the spectral PCs may underlie both hyperproliferative thickening and loss of orderly epithelial structure. The ability to link molecular trajectories with quantitative OCT imaging metrics highlights the integrated nature of morphological and biochemical remodeling during carcinogenesis.

### AI-driven multi-modal RS-OCT data fusion and classification

We then developed a convolutional neural network (CNN)-based AI framework to more efficiently extract, fuse and classify multi-modal RS and OCT data (Fig. 4a), effectively integrating the complementary strengths of both modalities. In our architecture, each modality was initially processed independently through a dedicated series of convolutional and maxpooling layers, allowing the network to learn hierarchical, modality-specific feature representations. The extracted features from the RS and OCT branches were then concatenated and passed through a set of shared fully connected convolutional layers to perform feature fusion and joint representation learning. This fusion strategy enabled the model to learn synergistic patterns across compositional and structural domains. Probabilistic classification to normal group and dysplasia/cancer group showed overall aggregated accuracy of 95.8% ± 3.4% (95% CI) (93.9% sensitivity and 97.8% specificity) for RS-OCT (Fig. 4b-c). By deactivating each modality, this enabled us to benchmark RS and OCT individually (Supplementary information Fig. S8). Their integration translated into a synergistic improvement in diagnostic accuracy.

**Fig. 4:**
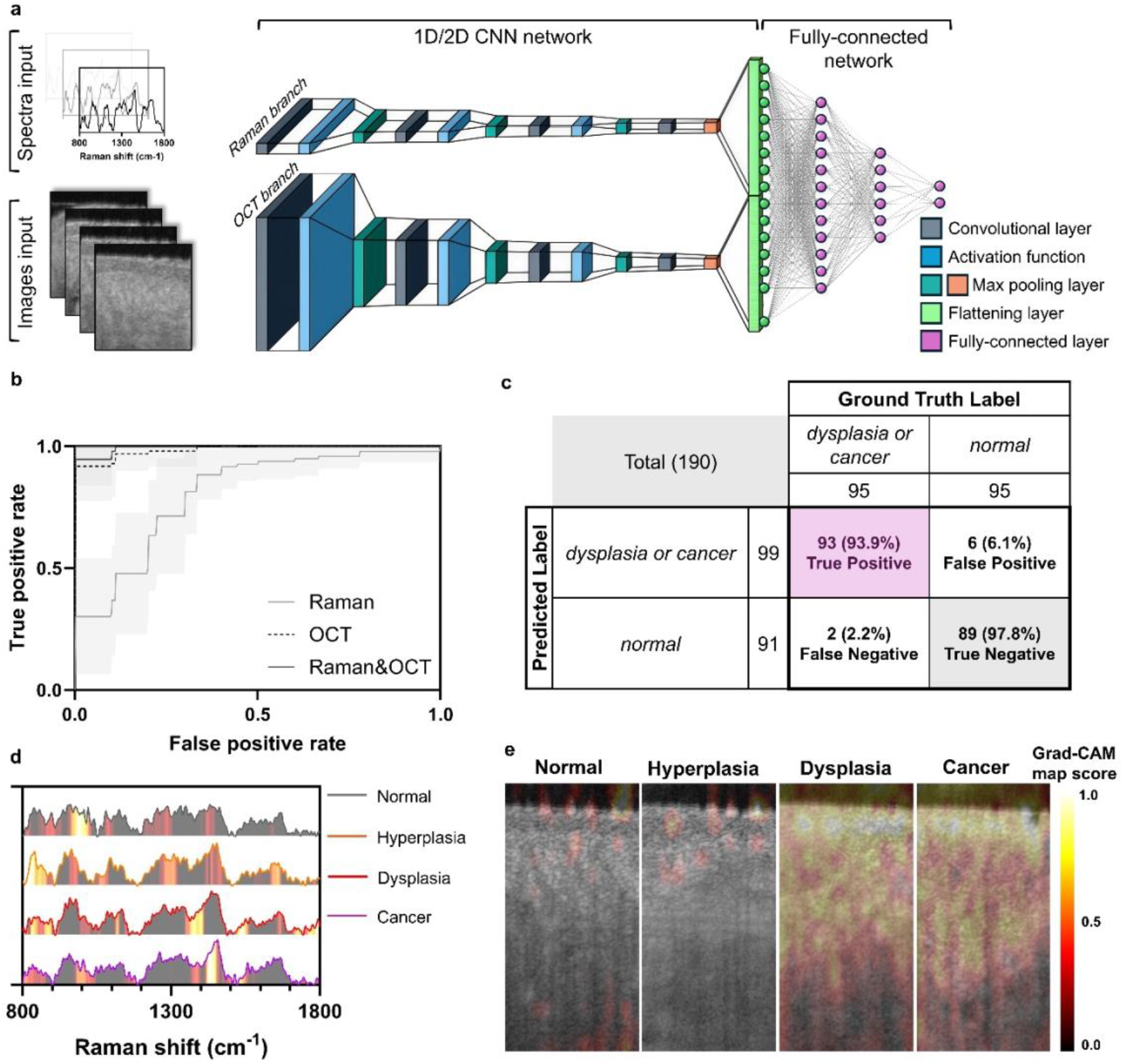
AI-driven multi-modal RS-OCT data fusion and classification. **a**, Deep learning architecture for temporal fusion of Raman spectroscopy (RS) and optical coherence tomography (OCT) data for disease classification. A custom-designed two-branch multimodal convolutional neural network (CNN) was developed to independently process spectral and imaging modalities before feature fusion. The RS branch comprises a 1D convolutional stack with 5 layers (32 to 128 filters) using ReLU activations and a kernel size of 5, optimised to extract spectral features across the Raman shift domain. This is followed by a 1D global average pooling layer to reduce dimensionality and capture global spectral patterns. The OCT branch employs a 2D convolutional backbone to extract spatial features from OCT B-scans, capturing epithelial morphology and microstructural changes. Features from both branches are concatenated and passed through fully connected layers for joint representation learning, followed by a softmax output layer for classification. **b**, Receiver operating characteristics curves (ROC) for differentiating between disease states using the multimodal RS-OCT fusion model. The model was trained to distinguish dysplasia and cancer from control tissue samples. **c**, Confusion matrix for classification based on fused RS and OCT data, illustrating the model’s accuracy and misclassification patterns. **d**, Grad-CAM heatmaps highlighting discriminative spectral features in Raman spectra for control, hyperplasia, dysplasia, and cancer samples. **e**, Grad-CAM visualisations revealing key diagnostic regions in OCT images across control, hyperplasia, dysplasia, and cancer.

We interrogated the decision-making process of our AI framework to ensure biological plausibility. Gradient-weighted Class Activation Mapping (Grad-CAM) was used to visualise the regions within the RS and OCT feature space that most strongly influenced individual classification outcomes (Fig. 4d-e). Exemplar Grad-CAM overlays were generated for correctly classified samples, revealing clear and modality-specific activation patterns. For RS inputs, Grad-CAM showed broad, distributed activations across the Raman spectral range, indicating that the model did not rely on a single peak or narrow feature but instead integrates information from the entire biomolecular fingerprint (Fig. 4d). For OCT inputs, Grad-CAM localised the most salient features within the epithelium across different tissue states. Interestingly, in control and hyperplastic tissue, attention was focused on epithelial structures, in particular normal shaped papillae in accordance with histology (Fig 2b) with minor emphasis along the basal layer. In contrast, dysplastic and cancerous regions showed strong Grad-CAM signals at structurally abnormal areas, particularly in the underlying stroma, consistent with sites of early pathological remodeling and tumour invasion (Fig. 4e). Therefore, the network leverages tissue microarchitecture and boundary integrity, key hallmarks used by pathologists, to drive its classification, effectively mimicking human diagnostic reasoning. Together, these findings demonstrate that the network learns representations across modalities, structural cues from OCT and molecular signatures from Raman spectroscopy, with high-intensity Grad-CAM regions aligning with biologically and diagnostically meaningful features. This multi-modal RS-OCT framework underscores its potential as a powerful tool for tissue classification and diagnostic decision support. The fusion allows for contextualised interpretation, linking molecular alterations to their corresponding architectural changes *in vivo*.

## Discussion

Current diagnostic methods for head and neck cancer primarily utilise white-light endoscopy and biopsy, which provide only intermittent snapshots rather than continuous monitoring. This approach misses the evolving molecular and microscopic changes in tumours over time, making early detection of progression from dysplasia to cancer challenging and potentially delaying crucial treatment decisions. Despite advances in widefield endoscopic imaging, *in vivo* longitudinal assessment of tissue at both molecular and microstructural levels remains a critical unmet need in oncology.

We presented a fully integrated dual-modality RS-OCT endoscopic platform that enables real-time, co-registered microstructural and biomolecular characterisation. This approach allowed dynamic monitoring of tissue changes during carcinogenesis. By leveraging a symmetric fibre geometry and a custom spliced-optic configuration, we achieved precise spatial and temporal co-registration of Raman and OCT signals, ensuring that biomolecular and architectural information is captured from the same tissue location effectively creating a dynamic snapshot of the tissue state. This opens new avenues for real-time, *in vivo* tracking of tumour evolution and response to therapy, with potential to fundamentally shift how early cancer is detected and monitored.

Applying this platform in a longitudinal 4NQO-induced oral carcinogenesis mouse model, we were able to temporally monitor disease progression from hyperplasia to dysplasia and eventually neoplasia. OCT imaging captured progressive epithelial thickening and loss of tissue stratification, while RS revealed molecular signatures indicative of increased nucleic acid and protein content. The combination of RS and OCT thus enables dynamic tracking of neoplastic transformation, allowing us to not only detect cancerous changes but also to observe the transitional states that precede malignancy. By longitudinally tracking these changes, the platform facilitates a deeper understanding of tumour evolution and heterogeneity, with RS effectively identifying molecular alterations and OCT robustly assessing lesion invasion.

The deep learning fusion model we developed enabled multi-modal representation learning, leveraging the microstructural and biomolecular contrast of each modality to enhance diagnostic classification. Grad-CAM visualisations consistently emphasised structurally relevant tissue regions while excluding common imaging artifacts such as speckle noise or motion distortions. This indicates that the model is not relying on acquisition-related artifacts but instead is focusing on meaningful tissue architecture across multiple depths, aligning with established histopathological principles where subsurface changes are critical for identifying dysplasia and cancer. A similar interpretability approach was applied to our RS branch, where importance mapping across the spectral domain revealed that the classifier prioritised specific Raman shifts corresponding to biologically specific molecular peaks such as 850, 1301 and 1650 cm^-1^ associated with collagen, lipids and proteins, respectively. These findings suggest that the model captures clinically relevant biochemical alterations associated with disease progression. Taken together, these explainability analyses reinforce that the deep learning models are leveraging biologically and clinically interpretable features, structural in OCT and biochemical in Raman spectroscopy, to make accurate predictions. This not only increases confidence in the model’s reliability but also provides a bridge between AI-driven insights and established diagnostic knowledge.

While the RS signal spatially is primarily derived from the region imaged by OCT, the Raman collection area extends slightly beyond the OCT B-scan field of view due to the wider spatial sampling of the collection fibres. Nevertheless, given the ∼1 mm axial imaging depth of the OCT and the spatial overlap in the probe design, the majority of the Raman signal corresponds to the region visualised by OCT. Future iterations of the platform may incorporate circular configuration with spiral or Lissajous scanning mechanisms to enable full en-face 3D OCT acquisition, thereby improving spatial co-registration with RS.

From a broader perspective, this work exemplifies how integrating advanced complementary optical modalities with AI can fundamentally reshape our diagnostic capabilities. Its compatibility with existing clinical endoscopes opens avenues not only for oncology, but also for inflammatory and degenerative diseases across the gastrointestinal tract where tissue morphological and molecular changes precede clinical symptoms. Looking ahead, we envision the integration of this dual-modality system with additional contrast mechanisms such as photoacoustic imaging, multi-photon imaging or fluorescence lifetime imaging to capture microvascular, functional and metabolic cues in parallel under widefield guidance^21^ or correlation with *ex vivo* spatial biology^22^. Hence combining these advances with real-time AI-driven analysis could yield a comprehensive *in vivo* biopsy. Further pragmatic developments could leverage AI enhanced imaging, wherein OCT image reconstruction with <100 ms RS clinical acquisition will be possible^23,24^. As optical technologies converge with computational advances, multi-modal endoscopy like RS-OCT may ultimately redefine the future of diagnostic medicine by enabling earlier, deeper, and more accurate temporal insight into disease state transitions and trajectory.

## Conclusion

We presented a compact, unified endoscopic platform integrating Raman spectroscopy and OCT within a miniaturised forward-viewing fibre-optic probe, enabling real-time, co-registered molecular and structural imaging *in vivo*. Combining subsecond RS acquisition with video-rate OCT and distal scanning, this system delivers unprecedented spatial and biomolecular resolution within a single optical field. Applied longitudinally in a head and neck carcinogenesis mouse model, it dynamically captured epithelial remodeling alongside molecular alterations driving malignant progression. This multimodal strategy transcends the limitations of individual techniques, providing a comprehensive, quantitative assessment of disease evolution. By fusing morphology with intrinsic molecular contrast, our approach pioneers label-free, real-time diagnostics and can revolutionise early cancer detection and guide precision interventions during clinical endoscopy.

## Materials and Methods

### RS-OCT instrumentation and fibre-optic probe fabrication

The RS-OCT instrumentation (Fig. 1a) was based on an (2D) spectral domain OCT scanning platform and an inhouse developed Raman system. For OCT we used a 1325 nm superluminescent light emitting diode (SLED) (SLD1325, Thorlabs, Inc: 1325 nm/ 10 mW/ 100 nm), a short-wave infrared Cobra 1300 spectrometer (C1300-1310/150, Wasatch Photonics, 147 kHz max line rate) and a fibre polarisation controller (FPC032, Thorlabs, Inc). The spectrometer was interfaced to a Camera Link image acquisition device (PCIe-1433, National Instruments Corp.). A visible laser (488 nm Toptica Ibeam Smart) was coupled in the same light path of the OCT to identify the location of the OCT output beam. The acquisition of OCT frames was synchronised to the beam scanning system using an analog output device (USB-6343, National Instruments). We used a 200 μm fibre (FG200LEA, Thorlabs, Inc.) coupled 785 nm diode laser for RS (Cleanlaze 500 mW, IPS photonics). RS system comprised a high throughput spectrometer (Acton LS785, Teledyne) with a thermoelectric cooled deep depletion charged coupled device (CCD) (Pixis 400×1340, Teledyne). The spectrometer had no slit so that the linear array of input fibre array (FG105LCA, Thorlabs, Inc.) acted as a slit.

The fibre-optic probe was constructed inhouse in a dedicated facility using motorised precision (< 5 μm) assembly (Fig. 1a-f, Supplementary Fig. S1). The fibre-probe employed a layered fibre arrangement resembling a sandwich, with two RS collection/excitation fibre arrays positioned on either side of a centrally aligned single-mode fibre (SMF-28 Ultra, Corning Inc.) dedicated to OCT. These RS fibres were angled slightly inward toward the optical axis. This configuration facilitated OCT B-scanning through a central slit, achieving a 1 mm field of view and delivering axial and transverse resolutions of 10 μm and ∼25 μm, respectively (Fig. 1d). The symmetrical layout preserved a fixed spatial relationship between the structural (OCT) and molecular (RS) sensing elements, enabling accurate alignment of morphological structures with their corresponding biochemical markers. Since RS is incompatible with distal quartz lenses, the OCT focusing optics were developed by splicing a NCF (FG125LA, Thorlabs, Inc.) and GIF (GIF625, Thorlabs, Inc.) fibre to the cantilever single mode fibre. We developed both a 3.4 mm and 2.2 mm diameter fibre probe.

To assemble the dual-modality probe, a piezo tube with four electrodes is first mounted to a stainless-steel tube (Supplementary Figure S1). The piezo tube was connected to a voltage source via four thin wires and secured to the stainless tube using a piezo holder. The motion of the free end of the 2.2 mm piezo tube (Physik Instrumente) was controllable by voltage. Next, ten 200 μm multimode low-OH silica fibres (FG200LEA, Thorlabs, Inc.) for Raman excitation and collection were cleaved flat and grouped into two linear fibre arrays, each containing five fibres. These were deposited with thin film bandpass/longpass filters (Shenzhen Photonstream Ltd). The linear fibre bundles were mounted onto a holder with a wedge design, forcing the two bundles to bend inward to increase the overlapping volume of excitation and collection with OCT single mode fibre. In the third step, a CaF_2_ was attached to the end facet of the linear fibre bundle holder. The holder was angled at the end facet to mitigate OCT back reflection. The CaF_2_ window sealed the probe and prevents contamination. Once the window was attached, an OCT fibre, consisting of a single-mode fibre and distal fibre optics, was gently inserted through the inner hole of the piezo tube using motorisation. The OCT fibre was positioned close to the window (∼150 μm) due to its limited working distance. Alignment of the OCT fibre was performed under a microscope for lateral positioning and using OCT imaging for axial positioning. Once aligned, the OCT fibre was secured to the free end of the piezo tube using epoxy adhesive. Finally, the proximal ends of the fibres were interfaced with the RS-OCT system. The clinically required OCT FOV of 1 mm is approximately equivalent or slight less that the size of biopsies. For the RS fibres, the central fibres of the two linear fibre bundles were used for excitation, where the Raman laser light is coupled in. The remaining Raman fibres were grouped into a round fibre bundle and connected to the spectrometer via an inline bandpass and longpass filters (BLP01-785R-25 & LL01-785-25, Semrock).

### RS-OCT real-time acquisition framework

We developed an acquisition framework in the Python environment. The software enables real-time RS and OCT data acquisition and analysis (Supplementary Fig. S2). This comprehensive real-time software package manages the fibre scan trajectory, generates synchronisation signals, reads the OCT and Raman spectra, and preprocesses the data. The fibre scanner and OCT spectrometer are hardware-synchronised to ensure precise image construction, while the Raman spectrometer is software-synchronised with the OCT spectrometer, allowing for temporal co-registration of the two modalities. Since the Raman spectrometer operates much more slowly than the OCT spectrometer, a multithreading design is employed to run the fast OCT data acquisition in parallel with the slower Raman acquisition. Raman data preprocessing includes simple dark current noise subtraction and intensity scaling. In contrast, OCT data preprocessing is more complex, involving background removal, k-domain linearisation, fast Fourier transformation (FFT), and normalisation. These OCT preprocessing tasks are offloaded to the GPU, enabling low-latency, video-rate rendering of the OCT images.

### Preprocessing and analysis of Raman data

The obtained spectra undergo wavelength calibration employing an atomic lamp (Ocean Optics HG-1). The background Raman spectrum of the system was subtracted from the tissue Raman spectra. To eliminate autofluorescence background, a custom fifth-order polynomial fit function^9^, constrained to the lower segment of the Raman spectrum was applied. Finally, the spectra were normalised to their integrated area to enabled relative abundance estimation and reduce absolute intensity fluctuations.

Principal component analysis (PCA) was then applied to reduce dimensionality and identify dominant sources of spectral variance. The number of principal components (PCs) retained for downstream analysis was determined based on the cumulative variance explained and the inspection of scree plots. PCs that collectively captured >99.29% of the total variance were initially considered. Individual components were further evaluated for their relevance to tissue pathology by inspecting loadings and their correlation with histopathological classifications. This approach ensured that retained PCs represented meaningful biochemical variation rather than noise.

### OCT image processing and quantitative morphological feature extraction

To extract morphometric parameters relevant to epithelial architecture and tissue organisation, we developed a quantitative OCT image analysis pipeline that processes B-scan images to derive two key metrics: epithelial thickness and an epithelial stratification/disorganisation index. These features were selected based on their clinical relevance to dysplasia and neoplastic progression in mucosal tissues. Raw OCT B-scans were first normalised to the fibre tip reflection (that appears as a horizontal bright line at the top of the supplementary OCT video) to exclude the fluctuation of light source. To mitigate sensitivity roll-off due to the finite-size of CCD pixel, depth-dependent normalisation was applied by dividing each A-line by a calibrated sensitivity curve. Following preprocessing, image segmentation was performed to delineate the epithelial surface and basal membrane. A semi-automated approach was employed to segment the epithelial layer:

1. The epithelial surface was identified as the first significant intensity rise along each A-line (first derivative Dijkstra’s-shortest-path-based edge detection).
2. The stroma surface was defined as the second significant intensity rise below the epithelial surface, reflecting the transition between the lamina propria and muscle. (ResNet-50 model trained by a manually labelled dataset, aided by an automated data labelling based on a first derivative edge detection)

Epithelial thickness was computed per A-line as the Euclidean distance between the epithelial surface and basal membrane along the axial direction. The final value for each OCT B-scan was derived by averaging thickness measurements across all valid A-lines (typically 256 per frame after down sampling). This provided a robust representation of overall epithelial thickening, a hallmark of dysplasia and malignancy.

To quantify epithelial architectural organisation, we defined a stratification index based on the variance in OCT intensity between the epithelium and stroma. For each A-line, the tissue depth was divided into two zones of equal thickness: a 21 μm lower epithelium region and a 21 μm upper stroma region. The zone thickness of 21 μm was chosen as a balance between two constraints: the minimum epithelial thickness in mouse tongues is approximately 42 μm, which sets the upper bound, and the OCT axial resolution is about 10 μm, which sets the lower bound. Selecting a midpoint value ensures that each zone is thick enough to capture relevant tissue structure while remaining within the resolution limits of the imaging system. We calculated the mean pixel intensities within these two zones for each A-line and computed a normalised stratification index as: *SI* = −20log (*I*_*e*_/*I*_*s*_), where *I*_*e*_ is the mean intensity of the lower epithelium and *I*_*s*_ is the mean intensity of the upper stroma. In well-organised (healthy) epithelium, stratification results in consistent reflectivity gradients between the basal layers and the muscle underneath, yielding predictable SI values. In contrast, dysplastic or disorganised epithelium exhibits disrupted polarity and reflectivity patterns, leading to altered SI ratios. All morphological features (mean epithelial thickness and stratification index) were computed for each imaging session per animal and temporally tracked across timepoints. These features were subsequently aligned with molecular (RS) features and endpoint histopathology labels for statistical analysis and multimodal classification.

### Temporal co-registration of OCT and RS

Analysing large volumes of multimodal data acquired at different temporal resolutions poses a significant challenge, especially in ensuring accurate alignment and integration across modalities. In our study, we combined RS, which provides one-dimensional spectral data at a rate of one spectrum per second, with OCT, which produces two-dimensional imaging at a high video frame rate of 46 frames per second. To align these temporally mismatched datasets, we implemented a straightforward temporal co-registration approach. Specifically, for each RS spectral acquisition (1 Hz), we randomly selected one of corresponding OCT frames captured within the same one-second interval. This method assumes that all OCT frames within that second are sufficiently similar for the purposes of co-registration. We chose this approach for its simplicity and practicality in exploratory data analysis. It allows efficient pairing of OCT and RS data without the need for complex synchronisation algorithms or hardware-based triggers.

### AI-based multimodal fusion and classification

To enable robust and generalisable morphomolecular classification from limited in vivo data, we implemented a deep learning framework for the fusion of RS and OCT data using a custom-designed multimodal CNN. The goal was to learn a shared representation that captures the synergistic diagnostic value of both biochemical and morphological features. The raw dataset consisted of n = 196 Raman spectra and n = 196 OCT B-scan frames, each co-registered and acquired from temporally tracked tissue sites in the oral mucosa of mice. All RS and OCT data were synchronised temporally and spatially via acquisition timestamps and scanner positional metadata. Since the mice study offered limited data of some groups, to mitigate overfitting and maximise generalisation, we applied imbalanced analysis extensive validation to avoid overfitting. The dataset was randomly stratified into training (72%), validation (18%), and test (10%) sets, ensuring that each set contained spectra and OCT frames. This was performed using a cross-validation strategy to provide statistical confidence estimates. Reported benchmarks (ROC, accuracy, sensitivity and specificity) were aggregated measures from the cross validation due to limited data in a 20 mice experiment.

We designed a two-branch multimodal CNN to independently process Raman and OCT data streams before joint fusion and classification. The Raman branch consisted of a 1D convolutional stack (5 layers, 32–128 filters, ReLU activations, kernel size = 5), followed by a 1D global average pooling layer. The OCT branch was implemented using a 2D convolutional backbone (based on a truncated ResNet-18 architecture), optimised for OCT’s texture-rich features. Both modality-specific feature embeddings were concatenated and passed through a series of fully connected layers (FC1: 64 nodes, FC2: 64 nodes) before a softmax output layer using a two-class weighted classification layer (control vs hyperplasia vs dysplasia vs cancer) with categorical cross-entropy loss to combat imbalanced datasets. Optimisation was performed using the Adam optimizer (learning rate = 1e–4, batch size = 512), with early stopping, choosing the most optimal model based on validation loss plateauing for 100 epochs. Model performance was evaluated only on the held-out test set using standard classification metrics, including accuracy, sensitivity, specificity, and area under the receiver operating characteristic curve (AUC). To assess the contribution of each modality, we deactivated either the RS or OCT branch, allowing us to quantify the added diagnostic value of multimodal fusion. All models were implemented in Matlab (v2023) and trained using a NVIDIA RTX A6000 GPU. Training took ∼few minutes per fold, and all results were aggregated over 10 cross-validation folds. The AI fusion framework is available at GitHub (www.github.com/bergholtlab/).

### Modal specific interpretability with Grad-CAM

To interpret the modality-specific contributions to the final classification, Grad-CAM was applied independently to each branch of the multimodal CNN. For the Raman branch, gradients of the output class score were computed with respect to the final 1D convolutional layer. These were used to generate a weighted activation map highlighting key spectral regions influencing the model’s decision. For the OCT branch, Grad-CAM was applied to the last convolutional block of the truncated ResNet-18 backbone. The resulting 2D heatmaps identified spatial regions within OCT images most relevant to the classification. By isolating Grad-CAM computation to each branch before fusion, we obtained interpretable, modality-specific visualisations of learned discriminative features.

### 4NQO murine carcinogenesis model

All animal work was approved locally at King’s College London (UK) and performed under a UK Government Home Office Project License (PP0313918). The animal experiment was designed with careful consideration of the 3Rs^25^. C57BL/6 mice were treated with 4-Nitroquinoline 1-oxide (4NQO) diluted to 100 μg/ml in drinking water for 16 weeks. 4NQO models are known to replicate the alterations caused by tobacco mutagens, progressing through a series of premalignant stages and exhibiting histological features that closely resemble human HNC^7^. Exposure to 4NQO leads to development of multi-focal dysplastic lesions in 100% of the treated mice. Of these, ∼50% will develop into invasive cancer within the 22 weeks. 4NQO-containing water was prepared and changed once a week for 16 weeks. During the experiments, the mice were maintained with regular mouse chow and water (with or without 4NQO) ad libitum (n=10 control and n=10 4NQO treated group). After that period, mice were given normal drinking water until the endpoint of 22 weeks or if mice losses > 15% of maximum weight. Over multiple time points treated, and control mice were sedated with inhaled isoflurane, and the oral cavities were screened for any visible lesions and RC-OCT data measurements.

### Histology

At the experimental endpoint, mice were euthanised in accordance with institutional animal care and use protocols. Whole tongues were carefully resected from the oral cavity using sterile surgical instruments. Care was taken to preserve both the tumour-bearing and surrounding non-tumorous mucosa for comprehensive histological evaluation. For frozen sections, tongue tissues were embedded in OCT (optimal cutting temperature compound), cryosectioned and stained with haematoxylin and eosin (H&E) by conventional methods. H&E slides were scanned with the NanoZoomer 2.0RS Digital Slide Scanner (Hamamatsu Photonics K.K., Japan) with 0.23 μm/pixel, 40× high resolution (Brightfield) mode.

### *In vivo* RS-OCT imaging of mice

For *in vivo* imaging, both treated and control mice were placed on a heating pad and sedated with inhaled isoflurane. The RS-OCT probe was gently positioned in direct contact with the mucosal surface at predefined anatomical sites within the tongue. To ensure consistent and reproducible sampling, probe placement was guided by anatomical landmarks and maintained using a micromanipulator arm. Raman spectra were acquired with an integration time of 1.0 s per location, while OCT imaging was performed continuously at a video-rate acquisition speed, enabling structural assessment in real time. A foot pedal was utilised to control whether to save the data to the disk. The optical power delivered to the tissue was 7 mW for the 1310 nm OCT channel and 60 mW for the 785 nm Raman excitation. To achieve temporal and spatial co-registration of Raman and OCT measurements, data acquisition was synchronised through a custom software interface that triggered simultaneous frame capture and spectral collection at each site, where the timestamp for each frame/spectrum acquisition was recorded. This ensured precise pairing of morphological and molecular data from the same tissue region across all imaging sessions. Repeated measurements were performed longitudinally at identical sites across timepoints (weeks 10–22) to enable morphomolecular tracking of carcinogenic progression *in vivo*.

## Data availability

The data that support the findings of this study are available from the corresponding author upon request.

## Code availability

The software used in this study is not publicly available. However, requests for code access for academic research purposes can be directed to the corresponding author.

## Acknowledgements

This work received funding from the MRC Impact Accelerator Award (MR/X502923/1) and the European Research Council (ERC) under the European Union’s Horizon 2020 research and innovation program (M.S.B., grant agreement No. 802778).

## Author Contributions

J.Q. designed the study, performed the experiments, interpreted, analysed the data, generated the figures and wrote the manuscript. P.G.B. designed and performed animal experiments, histology, contributed to scientific discussion, and data analysis. V.K contributed to experiments. J.G. performed histology. S.L contributed to histology. C.C. contributed with scientific discussion and data interpretation. R.J.C. contributed with scientific discussion and data interpretation. M.S.B. designed and supervised the study, interpreted and analysed the data, and wrote the manuscript.

