## Supplementary information for "Morphomolecular Longitudinal Endoscopy of Carcinogenesis"

### This PDF file includes:

#### Supplementary Figures

**Supplementary Fig. S1.**

**Supplementary Fig. S2.**

**Supplementary Fig. S3.**

**Supplementary Fig. S4.**

**Supplementary Fig. S5.**

**Supplementary Fig. S6.**

**Supplementary Fig. S7.**

**Supplementary Fig. S8.**

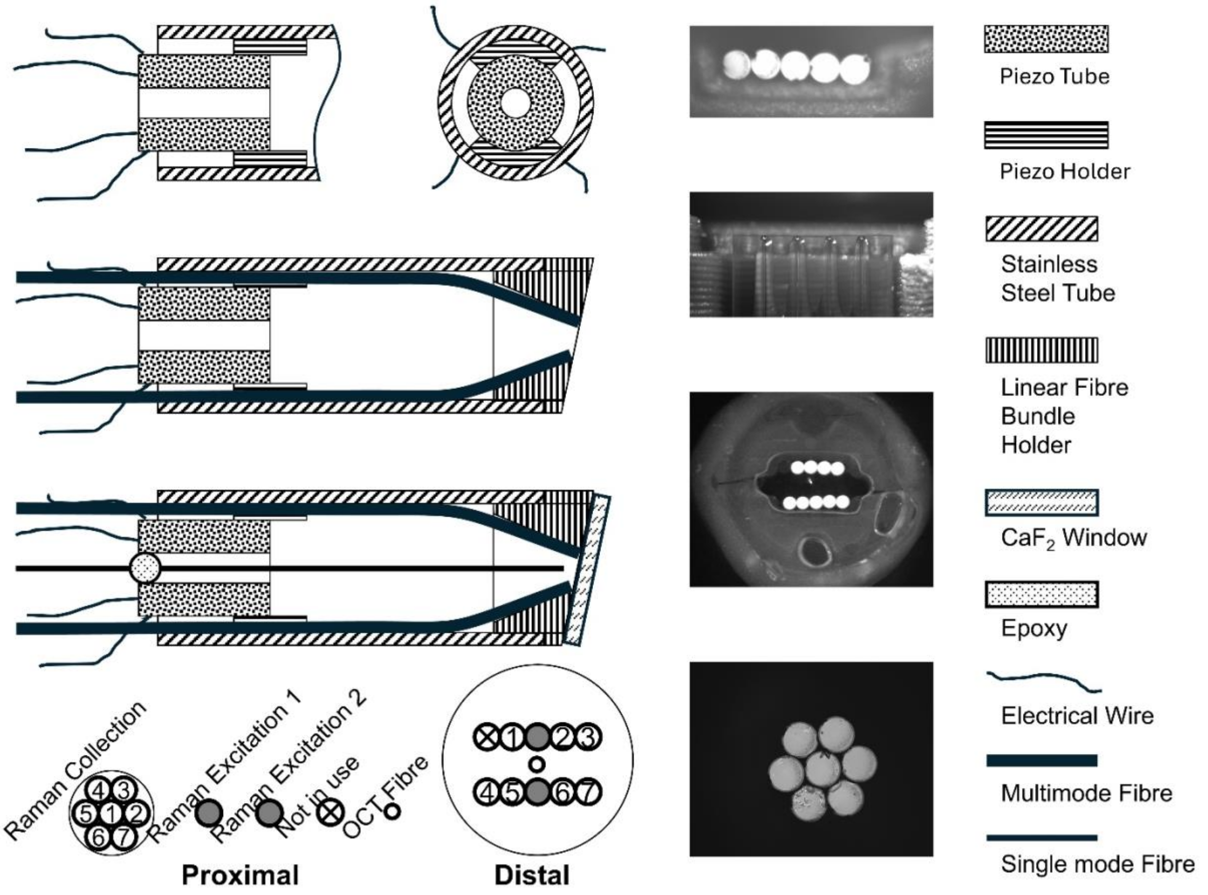


**Fig. S1: Design and assembly of the integrated fiber-optic RS-OCT probe.** Schematic diagrams (not to scale) show cross-sectional and longitudinal views of the distal and proximal ends. The probe integrates multiple fibers for co-localized optical coherence tomography (OCT) and Raman spectroscopy (RS). The distal tip contains a central single-mode fiber for OCT beam delivery, surrounded by six multimode fibers, two for RS excitation and four for RS collection, within a stainless-steel tube. A miniaturized, sandwich-like configuration positions two angled RS excitation/collection fiber arrays on either side of the central OCT fiber. Optical components are fixed with epoxy and a custom fiber-bundle holder, and a $CaF_2$ window seals the distal tip to provide optical access and biocompatibility. Microscopy images confirm precise fiber alignment. Labels indicate structural materials and optical elements used in fabrication.

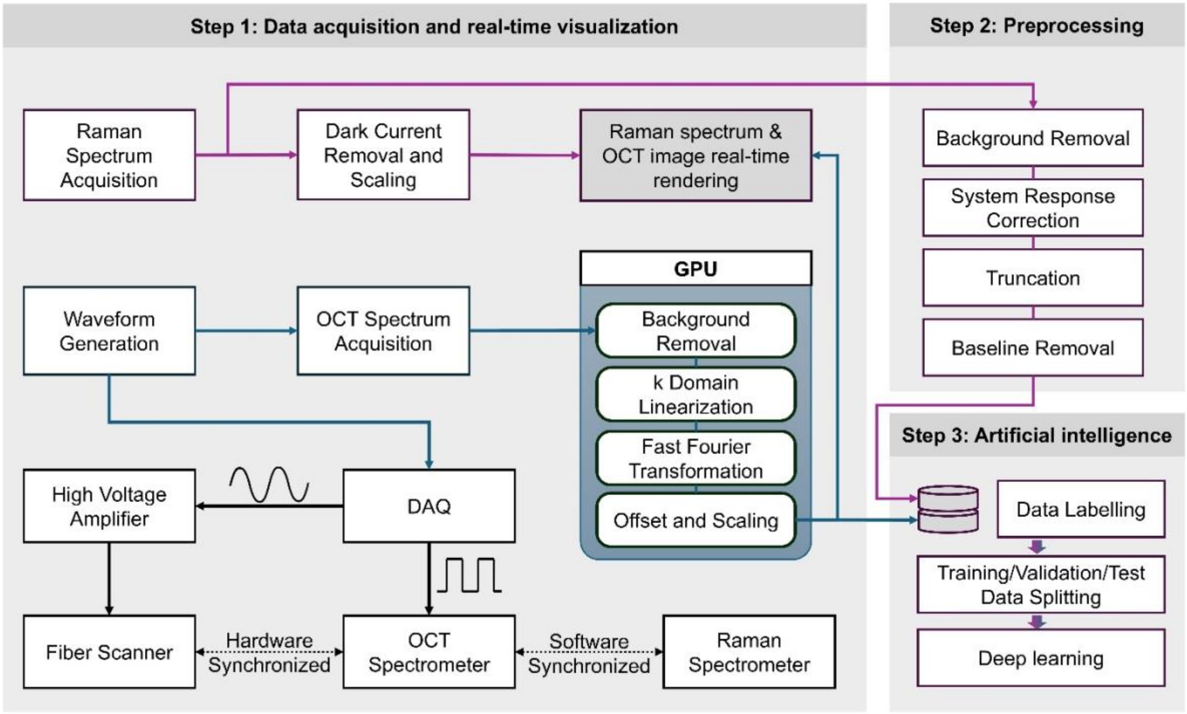


**Fig. S2: Schematic of the integrated data acquisition and processing framework for co-registered RS-OCT measurements.** A parallelized workflow enables simultaneous acquisition and processing of Raman spectroscopy (RS) and optical coherence tomography (OCT) data. OCT images are processed using a GPU-based pipeline comprising background removal, k-domain linearization, Fast Fourier Transform (FFT), and offset/scaling for high-resolution reconstruction. Raman spectra are processed on the CPU via background subtraction, autofluorescence removal by polynomial fitting, and vector normalization. Temporal co-registration is achieved by synchronizing each one-second RS acquisition with a randomly selected OCT frame from a 46-fps sequence, ensuring morphological (OCT) and molecular (RS) data correspond to the same tissue state.

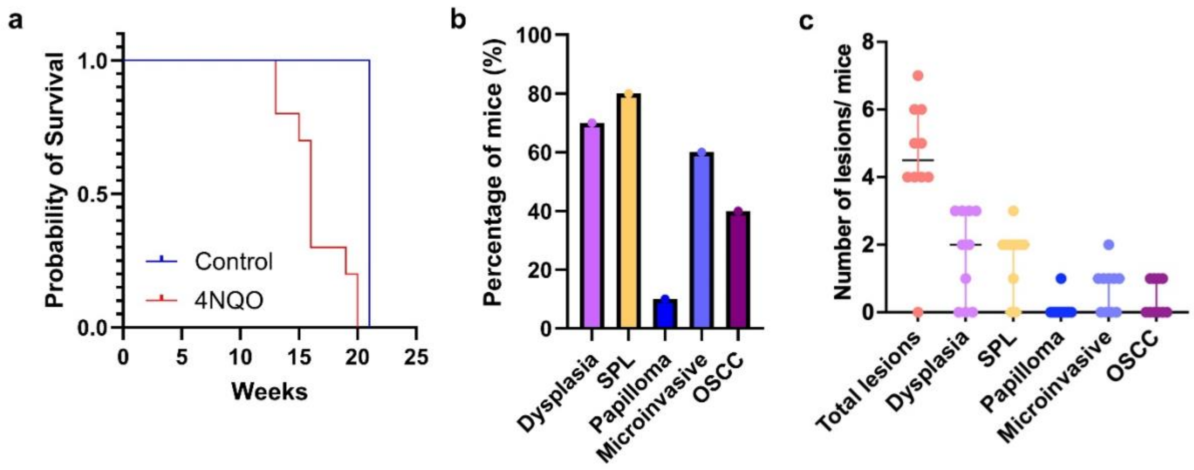


**Fig. S3: Survival plot and pathology outcome of mice in the 4NQO-induced oral carcinogenesis model. a,** Kaplan-Meier plots showing the probability of mice survival over the treatment and *in vivo* RS-OCT acquisition period in treated and control group (n=10 per group). Statistical test used: log rank test (*p<0.001)*. **b,** Endpoint histological outcome (percentage of mice with dysplasia, squamous papillary lesions (SPL), Papilloma, microinvasive and OSCC. **c,** Endpoint histological outcome (number of lesions per mice with dysplasia, SPL, Papilloma, microinvasive and OSCC.

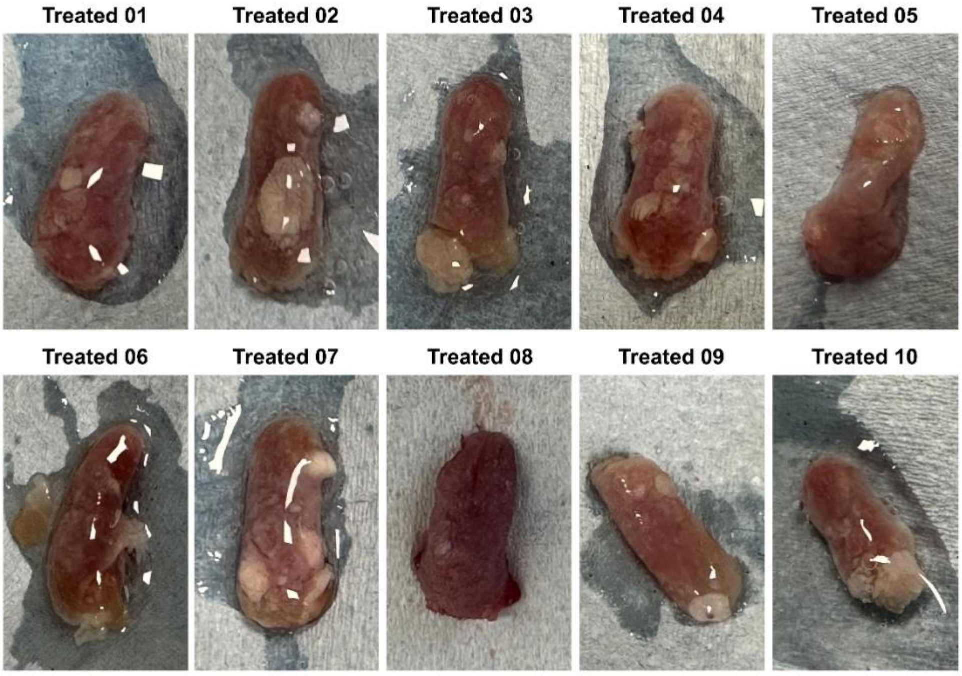


**Fig. S4. Macroscopic photos of resected tissues following 4NQO exposure.**

Representative images illustrate the heterogeneous presentation of oral lesions following 4NQO exposure. The gross morphological changes, visible as distinct lesions, correspond to a range of histological features, including, dysplasia, SPL, papilloma's and invasive OSCC.

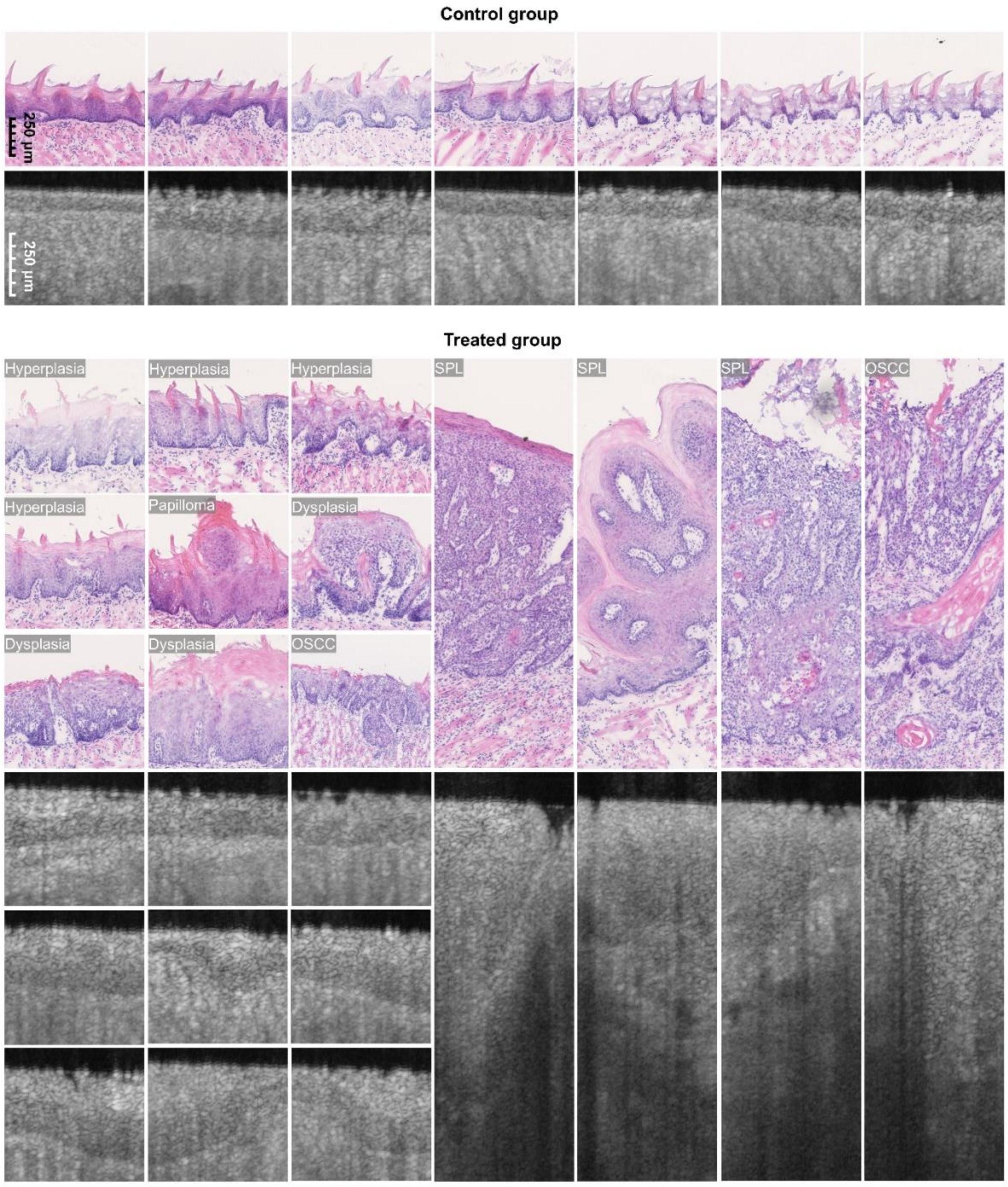


**Fig. S5: Additional representative H&E, OCT and RS data supporting the main findings.** Extended H&E-stained sections from control and 4NQO-treated tissue show oral lesions at various stages, including dysplasia of different grades, squamous papillary lesions (SPL), papilloma, microinvasive OSCC, and invasive OSCC. Additional in vivo OCT images from multiple animals and time points consistently reveal hallmark features of epithelial transformation, thickening, increased scattering, and loss of tissue organization.

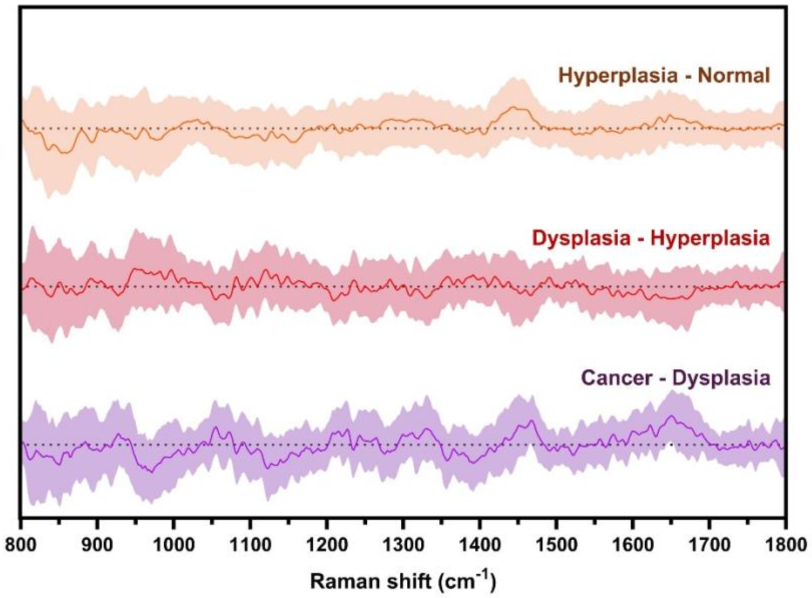


**Fig. S6: Raman difference spectra highlighting molecular changes during oral carcinogenesis.** Mean difference Raman spectra (±1 standard deviation) are shown for key pathological transitions in the 4NQO-induced murine model: control – hyperplasia, hyperplasia – dysplasia, and dysplasia – invasive cancer. Each spectrum depicts the mean normalized intensity difference between adjacent histological stages, revealing progressive biochemical alterations in tissue composition.

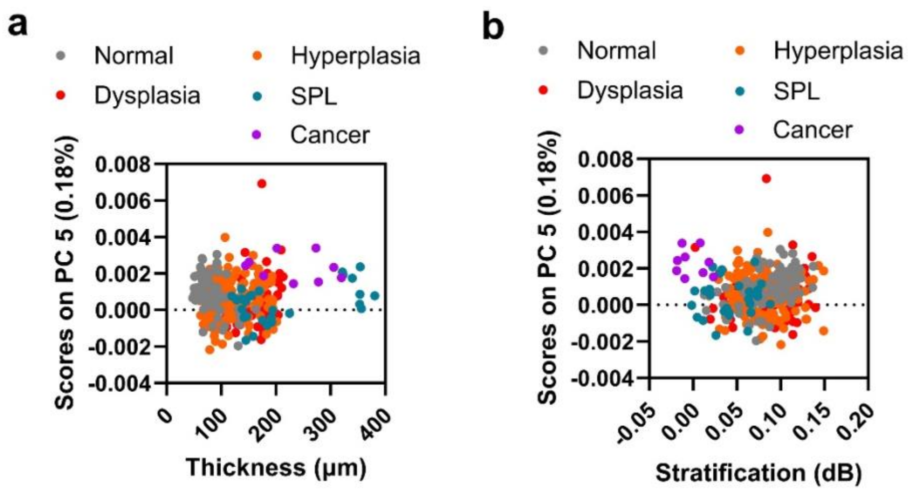


**Fig. S7: Morphomolecular analysis. a,** Principal component analysis (PCA) scores (PC5) plotted against epithelial thickness, illustrating the relationship between morphometric features and spectral variation across tissue samples. **b**, PCA scores (PC5) plotted against epithelial stratification for histopathologically confirmed control, hyperplasia, dysplasia, SPL, and cancerous tissues, highlighting how combined morphometric and molecular signatures differentiate pathological stages.

**Raman only**

| | | | Ground Truth Label | |
|---|---|---|---|---|
| | Total (190) | | *dysplasia or cancer* | *normal* |
| | | | 95 | 95 |
| **Predicted Label** | *dysplasia or cancer* | 102 | **74 (72.5%) True Positive** | **28 (27.5%) False Positive** |
| | *normal* | 88 | **21 (23.9%) False Negative** | **67 (76.1%) True Negative** |

**OCT only**

| | | | Ground Truth Label | |
|---|---|---|---|---|
| | Total (190) | | *dysplasia or cancer* | *normal* |
| | | | 95 | 95 |
| **Predicted Label** | *dysplasia or cancer* | 100 | **92 (92.0%) True Positive** | **8 (8%) False Positive** |
| | *normal* | 90 | **2 (2.2%) False Negative** | **88 (97.8%) True Negative** |

| | Sensitivity | Specificity | Accuracy |
|---|---|---|---|
| **Raman** | 72.5% | 76.1% | 74.2% |
| **OCT** | 92.0% | 97.8% | 94.7% |
| **Raman & OCT** | 93.9% | 97.8% | 95.8% |

**Fig. S8: Benchmarking performance of individual and fused modalities in the AI-enabled RS-OCT classification framework.** To evaluate the contribution of each imaging modality, we assessed classification performance using Raman spectroscopy (RS) and optical coherence tomography (OCT) independently. Classification models using only RS or only OCT inputs were trained with the same CNN architecture used in the fused model, with the corresponding modality branch deactivated.
